# Weakly-supervised Temporal Segmentation of Cell-cycle Stages with Center-cell Focus using Recurrent Neural Networks

**DOI:** 10.1101/2023.01.09.523193

**Authors:** Abin Jose, Rĳo Roy, Johannes Stegmaier

## Abstract

Training deep-learning models for biomedical images has always been a problem due to the lack of annotated data. Here we propose using a model and a training approach for the weakly-supervised temporal classification of cell-cycle stages during mitosis. Instead of using annotated data, by using an ordered set of classes called transcript, our proposed approach classifies the cell-cycle stages of cell video sequences. The network design helps to propagate information in time using Recurrent Neural Network and helps to focus the features on the center-cell. The algorithm is evaluated on four datasets from LiveCellMiner and has a performance close to the supervised approaches, which is impressive, considering that annotated data is not used in training.

## 1 Introduction

Mitosis is the process by which the cells rearrange and split into two identical daughter cells [1]. It is quite important to study and understand cell division to characterize pathological phenotypes in clinically relevant situations. The cell-cycle is divided into four phases: S, G2, Mitosis, and G1. The G1, S, and G2 phases are referred to as interphase. Mitosis is further classified according to the chromatin morphology into prophase, prometaphase, metaphase, anaphase, and telophase, where the pass to G1 is typically marked at cytoplasmic level by the cytokinetic process, which separates the cell bodies. Here we regard post-mitosis as the period where very gradual changes bring the compact anaphase chromatin to a decondensed G2-like mass in early G1.

In this work, we consider the classification of the cell-cycle stages into three different phases as in [2]: interphase, mitosis, and post-mitosis. Current state-of-the-art methods [2, 3] extract hand-engineered features, which are then classified using clustering methods. Zhong et al. [3] proposed an unsupervised clustering algorithm based on a temporally constrained combinatorial clustering (TC3) method. LiveCellMiner [2] additionally uses conventional convolutional networks to extract image features and uses a long short-term memory (LSTM) network to identify erroneous cell-cycle sequences. K-means clustering, after the feature extraction from fluorescence microscopy images, was proposed by Ferro et al. [4] for classification. Neural networks have strong feature extraction capabilities and is widely used in many computer vision applications [5–8] such as retrieval, hashing, segmentation etc. Many supervised models [9–13] are available for cell-cycle detection and classification. Recently, Jose et al. proposed a Recurrent Neural Network (RNN) based architecture [14] to incorporate time-related propagation of features for classification. However, this approach is supervised and requires careful annotations by a trained biologist. To overcome this problem, in this paper, we propose a weakly-supervised training scheme. The main advantages of this work are: 1) The network is end-to-end trainable and shallow. 2) The network focuses on the center-cell and occludes neighboring cells. 3) The classification head classifies the different states. 4) The training is weakly-supervised and needs only a transcript, a list of classes in order. For example, the transcript for a cell sequence is represented as : [interphase, mitosis, post-mitosis] as in the order of occurrence of classes, and it does not need dense image-to-image annotations.

## 2 Materials and methods

### Network architecture

For the temporal segmentation of the cell-cycle stages, we propose to use the same architecture as the one we proposed in [14], but the training method varies. The architecture of the model is shown in Fig. 1. The model architecture contains four main parts. 1) A backbone network to extract the features, 2) a time encoding network consisting of RNN layers to combine extracted features between the current frame and propagated features from the previous frame, 3) a focus network that helps to focus the extracted features towards the center-cell, and 4) a classification network to classify the cell-cycle stages. We used convolutional Gated Recurrent Unit (GRU) as the type of RNN in our model. The GRU connects the sequences temporally. This helps in the transfer of information between images in different time instances. The motivation to use convolutions in GRU was because our focus network reconstructs the image from high-level extracted features and focuses the features on the center-cell. The training of the model is explained in detail in the next section.

**Fig. 1.**
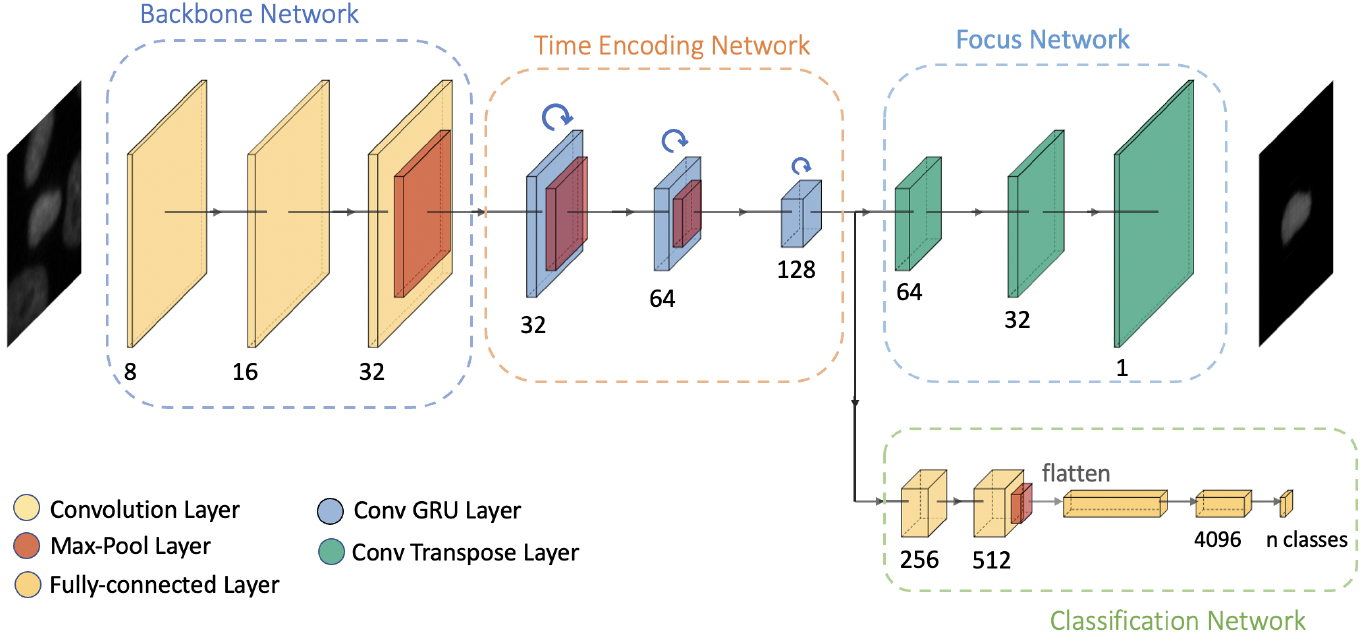
Illustration of our proposed architecture [14]. The model consists of two output networks, a focus network to reconstruct the center-cell and a classification network.

### Training details

The proposed model architecture contains RNN layers, which help the flow of information through time. Thus, the image at each time frame is dependent on the previous images from the sequence. Inspired by the Neural Network-Viterbi approach proposed in [15], our approach uses the Viterbi decoding method for the weakly-supervised training. For a given input sequence, given the order of occurrence of classes as a constraint, the weakly-supervised method finds the most probable set of labels. This set of labels contains the most probable class for each image in the input sequence under the given constraint. So, following training, the model gains the ability to recognize the classes for an input sequence given the classes’ order. As mentioned, our model architecture has two main output networks. The losses from these two networks are trained end-to-end to get the desired output.

### Focus network loss

The output of the focus network has a dimensionality equal to that of the input images. To reconstruct only the center-cell from the input image, the training calculates the loss between the predicted output and a masked image. This network aims to learn the features corresponding only to the center-cell in the input image and thus focus on the features of the center-cell. The masked image is obtained by using the annotated segmentation mask available with the datasets and setting all pixels that do not belong to the target cell to zero. The last layer of the focus network uses a sigmoid activation layer. The loss between the predicted and the expected input is calculated using binary cross-entropy (BCE) loss [16] and is given by:

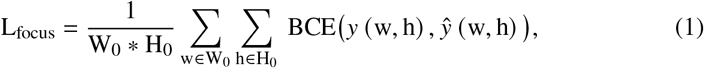

where *y* (w, h) and *ŷ* (w, h) are the input and predicted values of the network at pixel location (w, h). W_0_ and H_0_ are the input image dimensions. It is also possible to extract the segmentation mask of unseen images from this dataset using a simple threshold operation on the predicted output of the focus network.

### Classification network loss

The classification network treats the input as a sequence to determine the most probable path of classes. The computation of the most probable path and the loss associated with that path can be calculated only after getting the prediction outputs for all frames in a sequence. The classification network output is the posterior probability. The Viterbi algorithm [15] finds the most probable path using these posterior probabilities and a length model. Our approach used a class-dependent Poisson distribution as the length model as in [15]. Once the Viterbi algorithm finds the most probable path of labels, these labels are used to back-propagate through the network. A negative log-likelihood loss is used to train. Let the most probable path for an input sequence x^*T*^ found by the Viterbi algorithm be given by c_*n(t)*_. Then the weakly-supervised classification loss is given by:

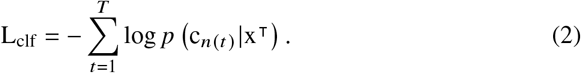

Both losses are computed for a sequence of data as input. A combined focus loss is obtained by summing the focus losses from each frame. The total loss is computed by a weighted sum of the combined focus loss and the weakly-supervised classification loss. A regularization parameter is used to combine them.

### 2.1 Datasets and experiments

#### Datsets

The datasets used for the evaluations are provided by LiveCellMiner [2]. The datasets contain microscopic image sequences of the mitosis process after the cell tracking. The datasets are acquired on human HeLa cells expressing H2B-mCherry transfected with indicated siRNA oligonucleotides in eight-well *µ*-slide chambers. It contains images that were taken three minutes apart. There are four datasets available from LiveCellMiner, namely LSM710 dataset, LSD1 dataset, RecQL4 dataset, and NikonXLight dataset. The former three datasets are captured using a confocal microscope and the latter is captured using a widefield microscope. Each contained cell is represented by a sequence of 90 frames, cropped from an image containing many cells to a resolution of 96 × 96 pixels with the target cell at the center. The dataset is annotated into three classes such as interphase class, mitosis class (prophase to early anaphase), and post-mitosis class (late anaphase to interphase). The post-mitosis class also contains cells that have grown into interphase (interphase recovery class) after cell division. Our approach is also able to identify the interphase recovery class, but we do not have a valid ground truth to evaluate instead, we qualitatively compared it with the findings from [2].

#### Experimental setup

For training the weakly-supervised model, a set of ordered labels called transcripts are used instead of the annotated data. The transcripts are simply the list of the classes in their order of occurrence. Our approach finds the number of frames in a sequence belonging to each class, given the transcript. Thus it helps to avoid the task of frame-to-frame annotation for training this model and the time involved for it. Given input sequences, the model trains both losses together with a regularization hyperparameter value of 0.01. The train-to-test data ratio is chosen as 0.85, and 10% of the test data is also used for validating the model. The batch size and the learning rate are selected as 4 and 0.01 respectively. The weakly-supervised classification loss is introduced every 20 iterations. The length of the transcript used is 4 considering the interphase recovery class as well. For evaluation, the interphase recovery class is considered a part of the post-mitosis class.

#### Evaluation criteria

The annotated data is available with the datasets for evaluation. We have calculated the frame-to-frame accuracy for each sequence and then averaged it. We also calculated the confusion matrix to find the correct classification and the misclassification rates. We compared the performance of our approach and compared the results obtained with the same model architecture trained using supervised methods [14] and the ResNet18 classification model.

## 3 Results

We plotted the label matrix in which the x-axis represents the length of each cell sequence, and the y-axis represents different sequences selected from the test dataset. Fig. 2 (a) shows the label matrix of the ground truth annotations compared with the label matrix of the predicted output on 50 test sequences of the LSM710 dataset. On closer inspection, it is evident that the predictions by the proposed model are similar to the user annotations. Fig. 2 (b) shows the first three principal components of the embedding space for the three classes of our proposed model. The embedding space contains the characteristics of the input data, in which the three cell-cycle states are clustered and move from blue to red to pink states and is clearly visible in the feature space. From the figure, it is clear that our proposed weakly-supervised method can cluster the features belonging to different classes without using fully-annotated data for training. Frame-to-frame accuracies can give a better quantitative analysis of the results. Tab. 1 shows the accuracies of the proposed model for the four datasets compared to the supervised learning approach [14] and also the ResNet18 classifier. Even though the proposed model has slightly lower (around 3-4%) accuracies when compared to the supervised methods, the results are impressive considering that dense ground truth annotations are not required for training. Fig. 3 shows the reconstructed image from the focus network for a given input image. The focus network reconstructs only the center-cell and thus helps the features learned in the embedding space to focus on the target cell. A segmentation mask of the target cell can be also predicted from the output of the focus network by using a simple intensity threshold.

**Tab. 1.** The frame-to-frame accuracy values of our proposed model in comparison with the supervised approaches [14] for the four different datasets.

| Model | Dataset |  |  |  |
| --- | --- | --- | --- | --- |
|  | LSM710 | LSD1 | RecQL4 | NikonXLight |
| Supervised [14] | 99.544 | 99.074 | 99.316 | 99.112 |
| ResNet18 [14] | 99.301 | 98.995 | 98.614 | 98.696 |
| **Ours** | 96.215 | 94.821 | 98.541 | 96.859 |

**Fig. 2.**
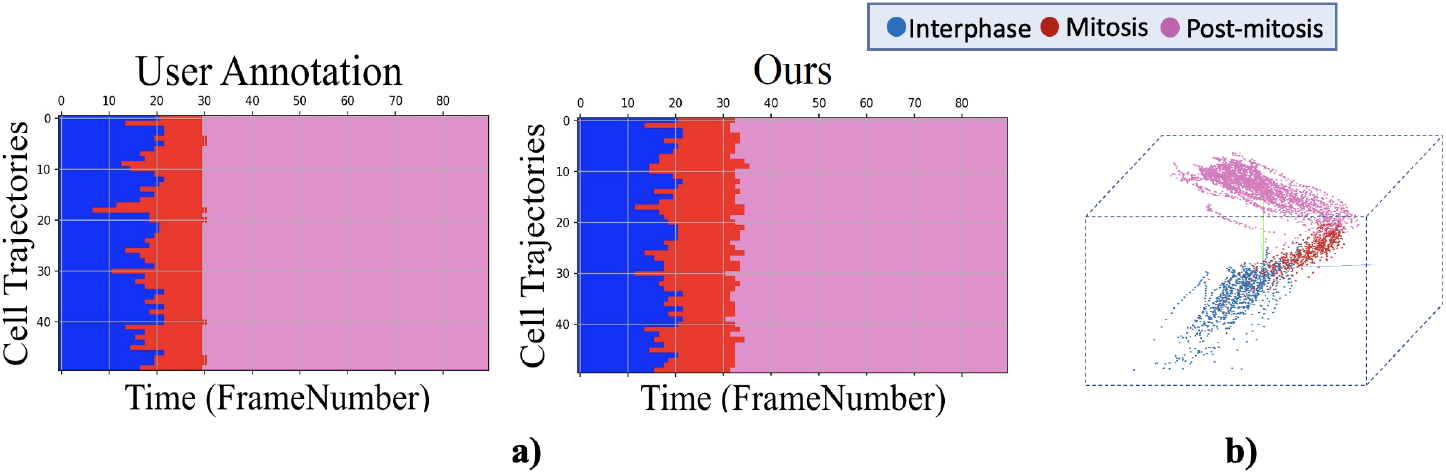
Illustration of the a) label matrix of the predicted output of our proposed approach compared with the user annotation, and b) first three principal components plot of the learned embedding space for the LSM710 dataset.

**Fig. 3.**
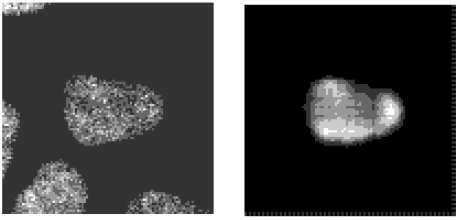
Input image (left), output of focus network (right).

Fig. 4 illustrates the confusion matrix of the proposed approach compared to the supervised approaches [14]. It is evident from the diagonal elements of the confusion matrix that the correct classification for three different classes is comparable to that with supervised methods. It is observed that the true positive rate for the mitosis class is higher than that of the other approaches. This indicates that the proposed weakly-supervised method can identify mitosis class images better than supervised approaches. For the other two classes, the miss-classification rate is higher for the proposed approach. It is also clear that confusion between unrelated classes does not occur as indicated by the zeros in the top right and the bottom left. Fig. 5 (a) demonstrates a plot from LiveCellMiner [2], which shows interphase recovery for the LSM710 dataset happens around 20 frames after mitosis. Fig. 5 (b) illustrates the label matrix of the proposed model when predicted into four classes. Even though there are no ground-truth annotations to evaluate an accurate measure, the qualitative results from the proposed model are in alignment with the findings from the LiveCellMiner.

**Fig. 4.**
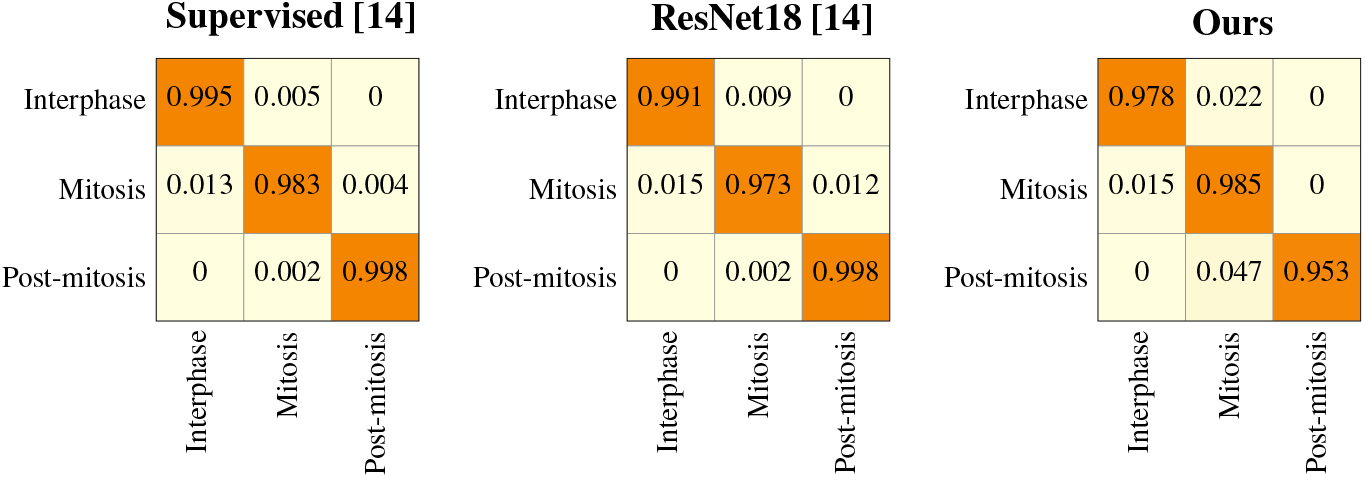
Normalized confusion matrices for the prediction on the LSM710 test dataset with our proposed model compared with supervised approaches [14].

**Fig. 5.**
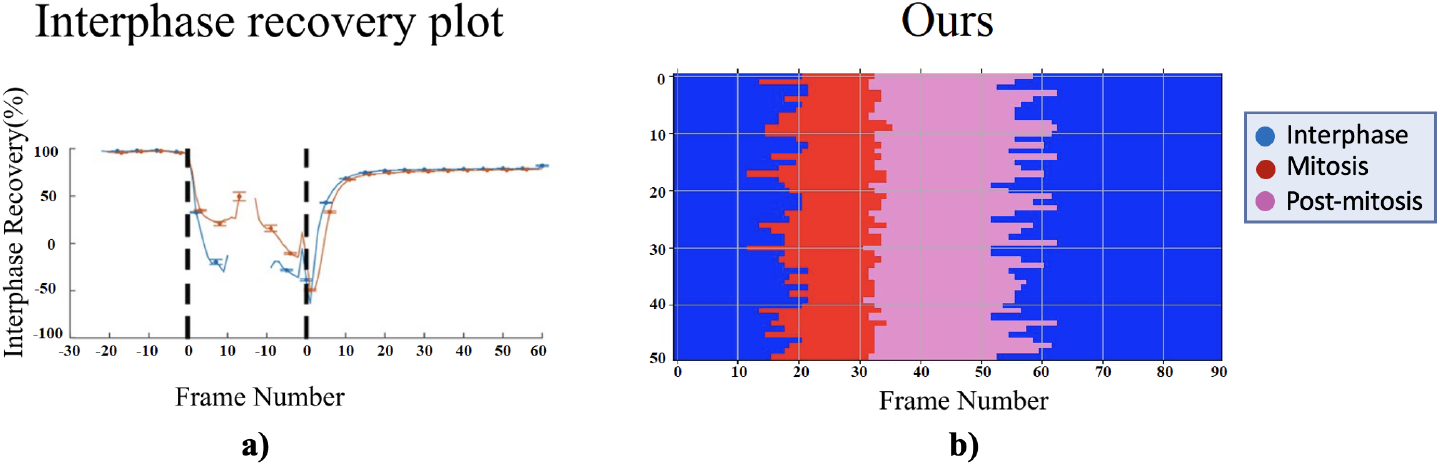
Interphase recovery class prediction. a) Plot from [2] illustrates that the interphase recovery occurs about 20 frames after mitosis. The indicated frame numbers are relative to two manually identified synchronization bars (left bar indicates interphase to prophase transition, right bar represents anaphase onset). The gap between the two transition time points helps to visualize aligned cell tracks irrespective of different durations of mitosis (Details in [2] Fig. S12). b) Our proposed weakly-supervised method when predicted into four classes predicted the recovered interphase frames (second set of blue after pink). The post-mitosis class (pink) covers a time span of 20-30 frames and is thus in line with panel a). Note that panel b) shows absolute frame numbers whereas relative frame numbers are used in panel a).

## 4 Discussion

We propose an automatic method for the identification of cell-cycle stages during mitosis. The proposed weakly-supervised approach only needs the transcript information which indicates the order of occurrence of cell-cycle states. Even though the classification accuracy is slightly lower compared to the supervised RNN model and ResNet18 classifier model, the proposed approach has the advantage that the user needs to provide only the order of occurrence of the states. This avoids the need for the generation of user annotation to train supervised models, which needs the expertise of a trained biologist. In the future, we would like to further extend the experiments to 3D cell datasets. Another interesting research direction is to use unsupervised training methods.

